# Modeling Time to Visual Insight in Mooney Image Recognition with a Chaotic Recurrent Neural Network

**DOI:** 10.64898/2026.01.12.699047

**Authors:** Misako Kimura, Yuuki Matsushita, Masayo Inoue, Shigeto Seno, Tsutomu Murata, Izumi Ohzawa, Toshio Yanagida, Kunihiko Kaneko, Kazufumi Hosoda

**Affiliations:** Center for Information and Neural Networks, Advanced ICT Research Institute, National Institute of Information and Communications Technology, 1-4 Yamadaoka, Suita, Osaka 565-0871 Japan; Department of Molecular Biochemistry, Graduate School of Science, University of Hyogo, 3-2-1 Kouto Kamigori, Ako, Hyogo 678-1297, Japan; Simons Centre for the Study of Living Machines, National Centre for Biological Sciences, Tata Institute of Fundamental Research, Kodigehalli, Bangalore 560065, India; Department of Basic Science, Faculty of Engineering, Kyushu Institute of Technology, 1-1 Sensui-cho, Tobata-ku, Kitakyushu 804-8550, Japan; Department of Bioinformatic Engineering, Graduate School of Information Science and Technology, Osaka University, 1-5 Yamadaoka, Suita, Osaka 565-0871 Japan; Department of Frontier Biosciences, Graduate School of Frontier Biosciences, Osaka University, 1-3 Yamadaoka, Suita, Osaka 565-0871 Japan; NEC Brain Inspired Computing Research Alliance Laboratories, Graduate School of Information Science and Technology, Osaka University, 1-5 Yamadaoka, Suita, Osaka 565-0871 Japan; Niels Bohr Institute, University of Copenhagen, Blegdmsvej 17, 2100 Copenhagen, Denmark; Department of Rehabilitation Science, Kobe University Graduate School of Health Sciences, Kobe 654-0142, Japan

**Author notes:** Corresponding author: Kazufumi Hosoda.

**Keywords:** Insight, Eureka, Aha, Recurrent neural networks, Artificial neural networks, Deep neural networks, Mooney image, Chaos, Optimization problems, Dynamical systems, High dimensional systems

## Abstract

Insight is often described as a sudden shift in or formation of a conceptual representation, enabling humans to restructure existing knowledge and solve problems beyond conventional analytical approaches. Although prior computational studies have modeled aspects of insight using deep neural networks (DNNs) or reinforcement learning, few have captured the dynamic emergence of insight through autonomous neural computation. Here, we present a neural network model that simulates the time required to reach visual insight in the Mooney image recognition task, a widely used paradigm for studying visual insight and perceptual reorganization. The model couples a DNN module for perceptual feature extraction with a recurrent neural network (RNN) that implements a chaotic search process for recognition. The RNN is formulated as a continuous-time dynamical system, autonomously explores internal states, and stabilizes when the missing visual features required for recognition are internally reconstructed. Using the same image set as in human psychophysical experiments, the model reproduces key statistical properties of human search times (STs), including (i) lognormal-like ST distributions across participants for each image, (ii) a proportional relationship between the log-scale mean and standard deviation estimated from lognormal fits across images, and (iii) discrete levels of the fitted log-scale mean across images (a proxy for image difficulty). Importantly, these properties emerge without assuming any lognormal distribution for participant-to-participant variability, whereas previous models reproduced similar signatures by positing lognormal-distributed individual differences. We further show that lognormal-like signatures can arise from exponential search dynamics when combined with both standard experimental preprocessing and finite observation windows, highlighting the need to distinguish generative mechanisms from measurement and analysis effects. Together, these results motivate a mechanistic link between chaotic neural dynamics and insight-related search and provide a computational framework for implementing insight in artificial systems.

## Introduction

Humans can arrive at innovative solutions by overcoming existing frameworks through insight. Since Gestalt psychology, insight has been understood as a problem-solving process driven not merely by trial-and-error learning but by a restructuring of the problem representation (Vitello and Salvi, 2023). Although various definitions exist, insight is often characterized as a sudden change in or formation of a concept or knowledge representation, typically associated with solving a difficult problem and accompanied by a pleasurable experience known as the “aha” or “eureka” effect (Kounios and Beeman, 2014). Insightful problem solving is thought to involve several stages, generally beginning when individuals encounter an impasse that cannot be resolved through analytical thinking, followed by a cognitive restructuring that leads to a discontinuous, emergent understanding (Kounios and Beeman, 2014). However, the computational and neurobiological mechanisms underlying such insight remain insufficiently understood.

From a computational perspective, several studies have attempted to identify the characteristics and key parameters of insight learning processes using reinforcement learning (Colin and Belpaeme, 2019; Harada, 2024; Chraibi Kaadoud et al., 2022), have observed phase transition-like phenomena resembling insight after extensive training of deep neural networks (DNNs) (Power et al., 2022; Clauw et al., 2024), and have demonstrated that adding random search strategies to DNNs can enable insightful problem solving (Trinh et al., 2024). In addition, coarse-grained models aligned with human experimental data have been proposed (Chao et al., 2025), and hypotheses grounded in neurocomputational modeling suggest that one-shot rapid neural plasticity mechanisms (Lansner et al., 2023; Wu and Maass, 2025; Hosoda et al., 2024) may play a crucial role in insight (Aru et al., 2023). However, no computational neuroscience model based on artificial neural networks (ANNs) has yet been developed that captures the entire insight process, including its emergent aspect involving the restructuring of internal representations. This may be because insight entails the restructuring of prior knowledge, making it difficult to define the necessary search space and evaluation functions.

Visual tasks have been extensively studied in both the brain and ANNs. For example, studies have investigated how neural clusters in higher layers represent visual features as prior knowledge (Tsunoda et al., 2001; Yamins et al., 2014). Therefore, compared to other domains, it is relatively easier to define the search space and evaluation functions for visual tasks. A widely adopted experimental paradigm for investigating visual insight involves the recognition of degraded images, such as Mooney images (not limited to faces in this context). Mooney images contain severely reduced visual information that typically fails to elicit recognition in the initial moments of viewing. However, with sustained observation, the concealed object may abruptly emerge into awareness, often accompanied by a characteristic ‘eureka’ experience (Mooney, 1957; Dolan et al., 1997; Tallon-Baudry and Bertrand, 1999; McKeeff and Tong, 2007; Giovannelli et al., 2010; Castelhano et al., 2013; Kizilirmak et al., 2016). From a neuroscientific perspective, this perceptual process appears to involve large-scale engagement across the brain that compensates for missing visual information via top-down mechanisms (Dolan et al., 1997; Hsieh et al., 2010; Gorlin et al., 2012; Hegdé and Kersten, 2010; Teufel et al., 2015; Chang et al., 2016; Martinsen et al., 2024; Jiang and He, 2006; Andrews and Schluppeck, 2004; Lu and Singer, 2023; González-García et al., 2018; Flounders et al., 2019; F. Imamoglu et al., 2012; Fatma Imamoglu et al., 2013; Casile et al., 2025; Murata et al., 2025). Recent work further suggests that one-shot, top-down influences in Mooney perception are primarily supported by higher-level visual cortex (e.g., HLVC/LO2) (Hachisuka et al., 2025). In addition, on the modeling side, incorporating attention into temporal sequence modules can yield more human-like one-shot learning effects in neural networks (Liu et al., 2022). However, the specific computational mechanisms underlying the search process that culminates in recognition after a prolonged period of non-recognition remain unknown.

The Mooney image recognition task is believed to involve a discontinuous process characteristic of insight (Murata et al., 2014; Hosoda et al., 2023; Smith and Kounios, 1996; Kounios and Beeman, 2014; Kounios et al., 2008; Salvi et al., 2016; Kizilirmak et al., 2016). One proposed mechanism is a coarse-grained model in which the human brain probabilistically activates the missing visual features of a concealed object in a degraded image, and recognition occurs when several of these features are simultaneously activated by chance (Murata et al., 2014). This model was motivated by three statistical properties observed in search time (ST) data, defined as the difference between the reaction time (RT) from stimulus onset to the recognition response for degraded images and the RT for the corresponding original color images: (i) for each image, the distribution of STs across participants followed a lognormal form, (ii) across images, the mean and standard deviation estimated from fitted lognormal ST distributions were proportional, with greater participant variability for more difficult images, and (iii) the distribution of image difficulty exhibited a discrete structure characterized by natural numbers.

Based on this coarse-grained framework, previous work constructed an ANN-based model (Hosoda et al., 2023) in which perceptual visual feature extraction was implemented using a DNN image classifier. The model used the same images as those employed in the human experiments and reproduced ST data with the same three statistical properties described above (Hosoda et al., 2023). However, in both the coarse-grained model and the ANN-based model, the cognitive search process leading to insight is fundamentally driven by random exploration and therefore is not embedded within the core neural computational architecture. Furthermore, although the lognormal property described in (i) has been attributed to a lognormal distribution of participant abilities, the origin of this lognormality remains unexplained.

In this study, we constructed a system that takes the same images used in human psychophysical experiments as input and outputs STs that exhibit the statistical properties observed in human data, with the cognitive search module also implemented using an ANN. Specifically, the perception module was implemented using a DNN image classifier, as in a previous study (Hosoda et al., 2023), whereas the cognitive search module was implemented as a dynamical system governed by time-evolving differential equations using a recurrent neural network (RNN). This RNN module adopts a model originally proposed to describe heuristic adaptation in biological systems such as intracellular and neural networks (Matsushita and Kaneko, 2023). The RNN performs exploration through deterministic chaotic dynamics and is designed to autonomously converge when the internally represented visual evidence becomes sufficient for recognition. Using this chaotic RNN, we reproduced the three hallmark statistical properties reported in human ST data. In contrast to previous accounts that rely on externally injected randomness, our approach generates these properties through intrinsic neural dynamics and does so without assuming lognormal variability in individual abilities. In addition, we show that the RT-to-ST transformation and the finite observation window used in psychophysical experiments can systematically shape fitted relationships in the log domain, highlighting the importance of explicitly modeling measurement and analysis steps when interpreting distributional signatures. Together, these findings establish a chaotic RNN as a mechanistic alternative to purely random-search explanations of visual insight and offer a route toward implementing insight-related recognition dynamics in artificial systems.

## Methods

### Input Image Dataset

The dataset consisted of 90 binary images and the corresponding 90 original color images. These images were previously used in psychophysical experiments (Murata et al., 2014). An example from this image set, a picture of a cut apple, is shown on the left side of Figure 1a.

**Fig. 1.**
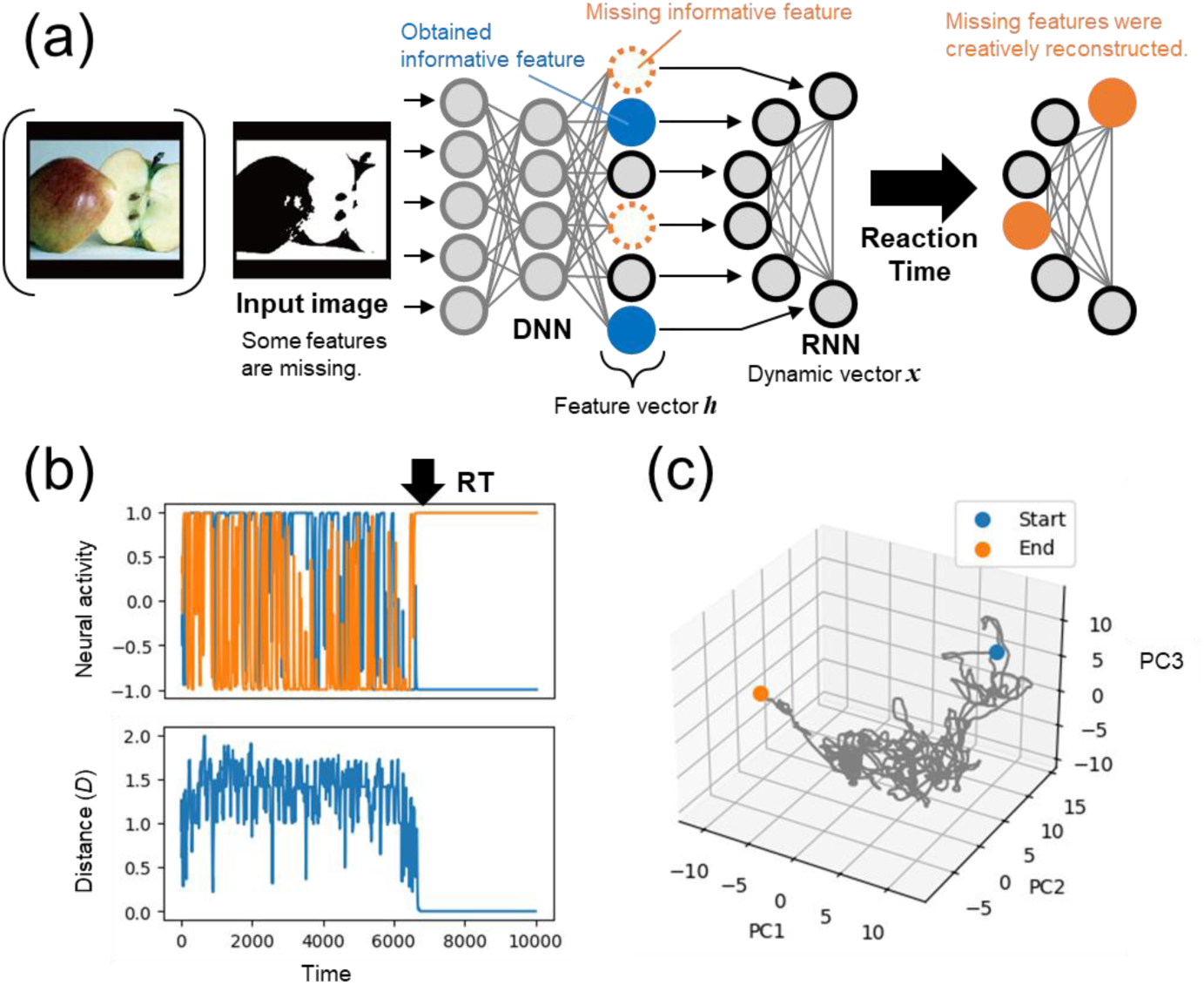
Overview of the model. **a** Schematic diagram of the model (see main text for details). **b** Example of time series data from a single trial. The top panel shows the activity of two representative neurons (out of the 768) in the RNN. The bottom panel shows the distance to the target. The arrow indicates the termination of the search, corresponding to the RT. **c** Trajectory of the RNN dynamics. The time series of the neuronal activity vector ***x*** is projected into three dimensions using PCA, and its evolution is shown as a gray line.

### Overall Model Architecture

We constructed a model that takes the same stimulus images used in human experiments as input and produces RTs as output. This model is implemented using two ANN modules: an image classifier DNN that extracts perceptual image features from the input images, and an RNN-based cognitive search module that compensates for missing features necessary for recognition (Figure 1a).

The use of the image classifier DNN as a perceptual feature extraction module was inspired by the concept of combination coding in neuroscience (Tanaka et al., 1991; Tsunoda et al., 2001) and by a previous study (Hosoda et al., 2023). Combination coding refers to the recognition of objects in higher visual cortical areas based on combinations of visual features represented by groups of neurons. Since this coding closely aligns with the mechanism of typical deep learning classifiers, we employed a standard image classification network to implement this functionality.

For degraded images, we assumed that the features available from the input are insufficient for immediate object recognition. Under this assumption, recognition occurs when the RNN-based cognitive search module compensates for the missing features through a process of trial and error, ultimately completing the assembly required for image recognition. The RNN used in this search module does not move optimally toward the correct answer, but rather evolves according to a predefined dynamical system. However, it spontaneously settles into a stable state once the correct answer is reached. This behavior allows us to define RT as the duration from the input of visual features into the RNN until the network autonomously stops.

It is important to note that the objective of our model in this study is not to classify degraded images per se, but to produce RTs. The model considers only the correct class and ignores incorrect classes. This reflects the design of the human experiment, in which the timer was stopped once participants reported recognition and restarted only if their response was incorrect. In this setup, incorrect responses were effectively treated as null events and excluded from RT measurement.

### Perceptual Feature Extraction Module

The feature extraction component follows the same configuration as in a previous study (Hosoda et al., 2023). We used ViT-B/32 (Vision Transformer) as the image classifier (Dosovitskiy et al., 2020; Morales). This model is pretrained on ImageNet with 1,000 classes (Deng et al., 2009) and is capable of extracting features sufficient for classifying those categories. However, since some of the 90 images used in our experiment did not belong to any of the original 1,000 classes, we employed Direct ONE-shot learning (DONE) (Hosoda et al., 2024) to add 15 new classes to the DNN, using three Creative Commons-licensed images per class obtained online. For each binary image, the “correct class” was determined by inputting the corresponding color image into the classifier and selecting, from among the top-1 or top-2 outputs, the class that was judged to broadly match the category of the image.

In the final hidden layer of the original DNN classifier, the classifier selects the class whose synaptic weight vector ***w****_i_* (corresponding to the *i*-th class) has the highest cosine similarity with the visual feature vector ***h*** (768-dimensional) extracted from the input. In this context, the weight vector ***w****_c_* for the correct class represents a typical pattern of visual features associated with inputs from that class. Based on this, we defined the necessary search space and evaluation function for restructuring prior knowledge in insight as follows.

Rather than considering all 768 features, we identified a subset of informative features for image recognition, following the concept of combination coding in neuroscience (Tanaka et al., 1991; Tsunoda et al., 2001). Specifically, we defined the “neurons that should fire” and the “neurons that should remain silent” based on the weight vector ***w****_c_*, selecting those with the top 5% and bottom 5% of weights, respectively. We then calculated *K* as the total number of such neurons in the feature vector ***h*** that were actually active (top 5%) or inactive (bottom 5%). The number of missing informative features (*N_M_*) was defined as *N_M_* = max (0, *K_thr_* - *K*), where *K_thr_* is the recognition threshold for *K*. From the informative features not counted in *K*, *N_M_* elements were randomly selected, and recognition was considered successful when these features were modified to match their corresponding states in ***w****_c_*. The threshold *K_thr_* was set to the minimum *K* value observed among the 90 color images (*K_thr_* = 10), and this same threshold was uniformly applied to all 90 binary images, since all participants in the human experiments were able to recognize every color image instantly.

### Cognitive Search Module

The search module takes as input the visual feature vector ***h*** extracted from the input image and employs an RNN consisting of a state vector ***x*** with 768 neurons, each corresponding to a neuron in the feature vector. The activation *x_i_* of the *i*-th neuron evolves over time according to the following equation:

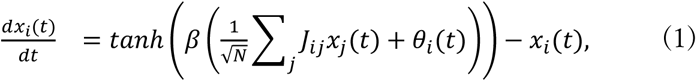

where *J_ij_* represents the synaptic strength from the *j*-th neuron to the *i*-th neuron in the connectivity matrix ***J***, and *θ_i_* denotes the excitability of the *i*-th neuron. The tanh function serves as a sigmoid, and following the prior work (Matsushita and Kaneko, 2023), we set *β* = 40 to approximate a binary on-off switch. When the sum of influences from other neurons and *θ_i_* is positive, *x_i_* quickly increases to 1, whereas when it is negative, *x_i_* rapidly decreases to -1. The excitability parameter *θ_i_*, which influences firing independently of interactions with other neurons, changes slowly over time according to:

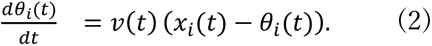

Equation (2) describes a first-order lag process in which *θ_i_* approaches *x_i_* at a rate determined by *v*. When *v* is large, a high *x_i_* leads to an increase in *θ_i_*, making *x_i_* more likely to remain active. In this way, the current state of *x_i_* is reinforced and the behavior becomes gradually stabilized. In this study, we define the rate *v* as the parameter that controls the convergence behavior of the module:

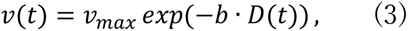

where *v_max_* is the maximum value of *v* and *b* controls the dependence of *v* on *D*(*t*). We used the default parameter values from the previous study (*v_max_* = 0.1 and *b* = 4 in Eq. (3)) (Matsushita and Kaneko, 2023) and confirmed that the main conclusions of this study remained unchanged when each of these parameters was varied by a factor of two. *D*(*t*) denotes the mean Euclidean distance between the current activations *x_m_* of the *N_M_* missing informative features and their target values *X_m_*, defined as:

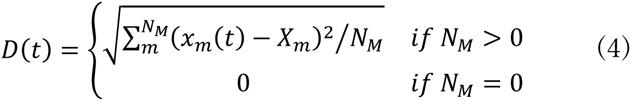

These target values *X_m_* are derived from the correct class weights ***w_c_***: neurons in the top 5% are assigned a value of 1 (firing), and those in the bottom 5% are assigned -1 (inactive). When all *N_M_* missing features have been successfully complemented with their target values, ***x*** stabilizes at the maximum rate *v_max_* (0.1). Nevertheless, since *v* is always less than 0.1, *θ_i_* evolves more slowly than *x_i_*, even at its maximum value.

For each degraded image, the temporal evolution of ***x*** was computed using the above model. The initial value of ***x*** was obtained by applying quantile normalization to the corresponding feature vector ***h*** using the standard normal distribution, followed by normalization such that the maximum value was 1 and the minimum was -1. All initial values of *θ_i_* were set to zero. The simulation was terminated when both *D* and the absolute change in all *x_i_* values became less than 10^-6^ and the elapsed time was defined as the RT. We simulated 100 virtual participants, each assigned a unique ***J*** matrix in which the synaptic weights were randomly selected from the set {-1, 0, 1} with equal probability.

### Conversion from reaction time to search time

In the previous human experiments (Murata et al., 2014), RT was defined as the interval from the onset of a degraded image to the recognition response. ST was estimated by subtracting the RT for the corresponding original color image, which was treated as an estimate of “non-search time.”

In the present model, we treat the time required for the RNN to reach a stable termination state as the model RT. This is because, by construction, the RNN requires a finite amount of time to terminate even when no search is performed (*N_M_* = 0). Accordingly, we define the model non-search time as the minimum termination time in the *N_M_* = 0 condition. Model ST is then computed by subtracting this non-search time from the model RT for conditions with *N_M_* ≥ 1. Images with *N_M_* = 0, which by definition involve no search process, are excluded from analyses of ST.

## Results and Discussion

### Model Behavior

Our model consists of two modules: a DNN for perceptual feature extraction and a chaotic RNN for cognitive search (Figure 1a; see Methods for details). The RNN search module does not optimize directly toward the correct answer; rather, the state variable ***x*** evolves according to a predefined dynamical rule (Eq. (1)), and its excitability ***θ*** changes over time following Eq. (2). While individual neurons do not know how to move toward recognition, they evolve according to the predefined dynamics and are influenced by a global measure of distance to recognition that is shared across the system. To illustrate the model’s behavior with a simple example, when *N_M_* is two, the system initially explores freely, but once those two features are filled, the entire network spontaneously stabilizes into a fixed pattern of activation and inactivation. This behavior is an intrinsic property of the model (see Methods).

Figures 1b and 1c illustrate an example trial, showing the temporal evolution of two selected neurons out of 768, the change in distance to the target, and the trajectory of the full RNN state in a principal component analysis (PCA) space. Unlike typical optimization methods, which begin with coarse adjustments and gradually refine the solution, our model does not follow such regularity. Instead, it exhibits behaviors such as large fluctuations occurring mid-process or abrupt transitions to the correct state, as shown in the example. This example indicates that the system does not exhibit consistent or periodic behavior even within a single trial, suggesting the presence of chaotic dynamics. Indeed, the maximum Lyapunov exponent was positive in all cases tested.

### Reproduction of Lognormal-Like Distributions Observed in Human Experiments

For each image, we converted the RTs of 100 simulated participants into STs and examined the cumulative frequency distribution of log_10_-transformed STs. The resulting distributions exhibited a lognormal-like shape, similar to those observed in human experiments (Figure 2a; normal probability plots yielded R² = 0.98, comparable to the previously reported human value of R² = 0.97, computed as the mean across 41 images with the shortest STs, for which 90% of participants reached recognition in the human experiments). Although there were slight mismatches between the simulation and the human experiments in scale, we used the default RNN parameter values from the previous work in this study (see Methods); tuning these parameters could improve the scale match. It is important to note that the previous modeling study (Hosoda et al., 2023) assumed a lognormal distribution of participant ability to account for the lognormality observed in RTs. In contrast, the current model makes no such assumption, yet still reproduces lognormal-like behavior.

**Fig. 2.**
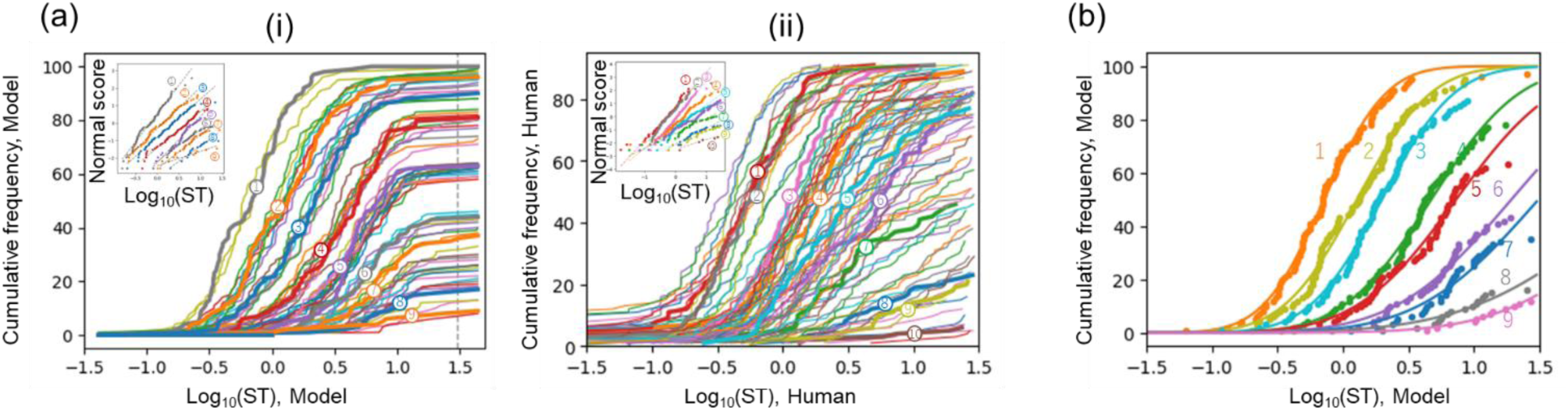
Cumulative distributions of STs. **a** Cumulative distributions across all images. (i) and (ii) show the results from the present simulations and the previous human experiments, respectively. Each line represents a single input image, and the vertical axis indicates the cumulative frequency of participants who had completed the search by each time point. Inset: normal probability plots of log10(ST) versus normal scores for representative stimuli; the solid lines indicate linear fits. **b** Lognormal distribution fitting. One representative example is shown for each *N_M_* value (indicated numerically).

Another characteristic of the model is that it tends to halt after a certain period of time, even when a solution has not yet been reached (Figure 2a). As a result of this premature termination, the cumulative distribution plateaus after a certain time point, reflecting simulated participants who fail to reach a solution within the simulated time window. In the human experiments, the upper limit was set to 30 s, and thus it remains unclear whether participants would have reached a solution if given more time. However, it is plausible that sustained effort becomes difficult over prolonged periods, and the empirical distributions do not rule out such a possibility. While such premature termination is a property of the model, it may point to a comparable feature in human cognitive processing.

To approximate the 30-s cutoff used in human experiments, we defined the time at which the last participant completed the search for the most difficult image as the time-on-task limit, yielding comparable success rates of 7% and 5% for model and human participants, respectively (Figure 2a, gray dashed line). Under this constraint, we fitted the data with a lognormal distribution, as illustrated in Figure 2b (examples from 9 images).

### Reproduction of Statistical Properties Observed in Human Experiments

As in the human experiments, we observed a proportional relationship between the mean and standard deviation (SD) obtained from the lognormal fits described above (Figure 3a). In addition, as also reported in the human experiments (Murata et al., 2014), the histogram of fitted mean values exhibited a discrete structure (Figure 3b), with zero-frequency bins separating clusters corresponding to *N_M_* = 1, 2, 3, and 4. This discreteness was preserved in additional simulations with an increased number of images (100 virtual images for each *N_M_* value from 1 to 9; Online Resource 1). Except for extremely difficult cases (*N_M_* ≥ 8), the spacing between clusters was approximately uniform, consistent with the human data, in which only the discreteness of the fitted values could be discussed due to the unobservability of *N_M_*.

**Fig. 3.**
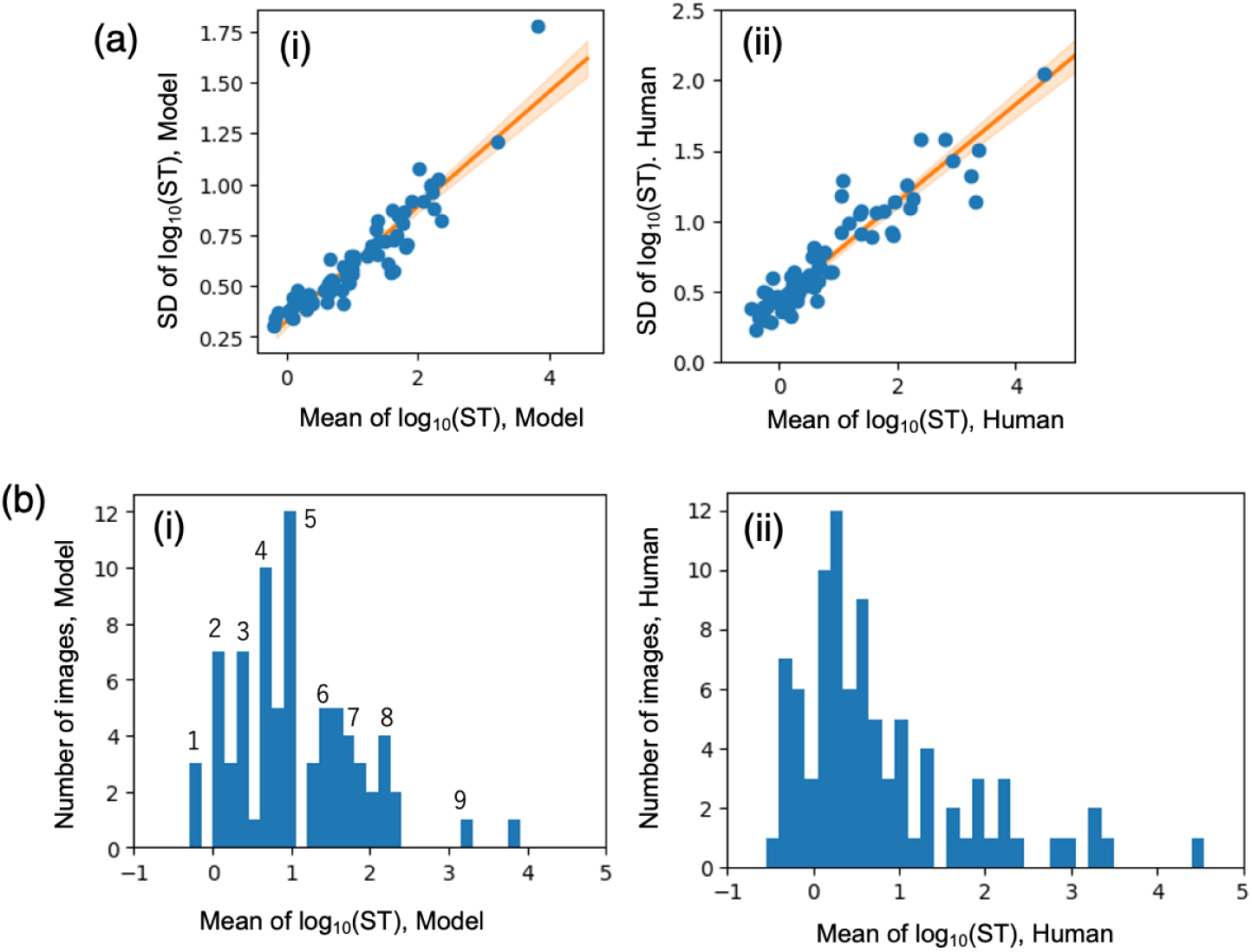
Statistical properties of fitted ST across images. (i) and (ii) show the results from the present simulations and the previous human experiments, respectively. **a** Relationship between the mean and SD of log-transformed STs across images. Each point represents one of the 90 images. The line shows the linear regression fit, and the shaded area indicates the 95% confidence interval (CI). The slope, intercept, and R² were 0.28, 0.33, and 0.88 for (i), and 0.35, 0.45, and 0.89 for (ii), respectively. **b** Histogram of the mean log-STs across images. Numbers indicate the corresponding *N_M_* values (available only for (i)).

The reason why log_10_(ST) exhibits an approximately evenly spaced structure is that the ST is defined as the time at which *N_M_* units simultaneously match the target pattern. For example, if the firing probability of each neuron is assumed to be constant at *p*, the expected time until *N_M_* neurons fire simultaneously is proportional to 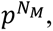 implying that ST scales multiplicatively with increasing *N_M_*. Such scaling behavior is consistent with results reported in the previous random-search model (Hosoda et al., 2023). The present study demonstrates that the same property can also be reproduced within a framework based on chaotic search dynamics.

For reference, since the input images used in the present model were identical to those used in the human experiments, we examined the relationship between model and human RTs. The correlation was weak, although it was statistically significant (Pearson’s *R* = 0.24, *p* = 0.022; Spearman’s *ρ* = 0.29, *p* = 0.006). As shown in Figure 3b, RT is largely determined by *N_M_*, and therefore this correlation primarily reflects the relationship between ViT-based feature extraction and human perception, rather than the influence of the RNN dynamics. These results suggest that the model does not yet match human performance, while offering a starting point for future quantitative comparisons to assess the remaining gap.

### Mechanistic origin of the observed mean-SD correlation

In the previous human study (Murata et al., 2014), the origin of the lognormal distribution was interpreted as reflecting lognormally distributed variability in participants’ abilities, whereas the dynamical properties of the search process itself were not fully elucidated. In contrast, the present study allows us to directly examine the statistical properties of the search process implemented in the model.

The chaotic search model used in this study, originally proposed as a generic form of biological optimization (Matsushita and Kaneko, 2023), incorporates both non-search time and premature convergence, as described above. As a result, the distribution of pure search time was obscured and was not explicitly characterized in the original study. Here, we analyze the distribution of pure search time.

To isolate the pure search dynamics, we first address premature convergence specific to this model, which arises from the exponential term in Eq. (3) implementing slow fixation. To remove this effect, we modified Eq. (3) by replacing the exponential term with a Boolean gating rule such that *v* is set to *v*_max_ when the distance becomes sufficiently small (*D* < 10_-0.5_); otherwise, *v* is set to zero. Under this modification, premature convergence was substantially suppressed (Figure 4a). When the effect of non-search time is negligible, namely for sufficiently large *N_M_* (e.g., *N_M_* ≥ 5), the process is dominated by chaotic searching. In this regime, the cumulative frequency distribution of search times is well approximated by an exponential distribution (Figure 4a), indicating mechanistic similarity to random search.

**Fig. 4.**
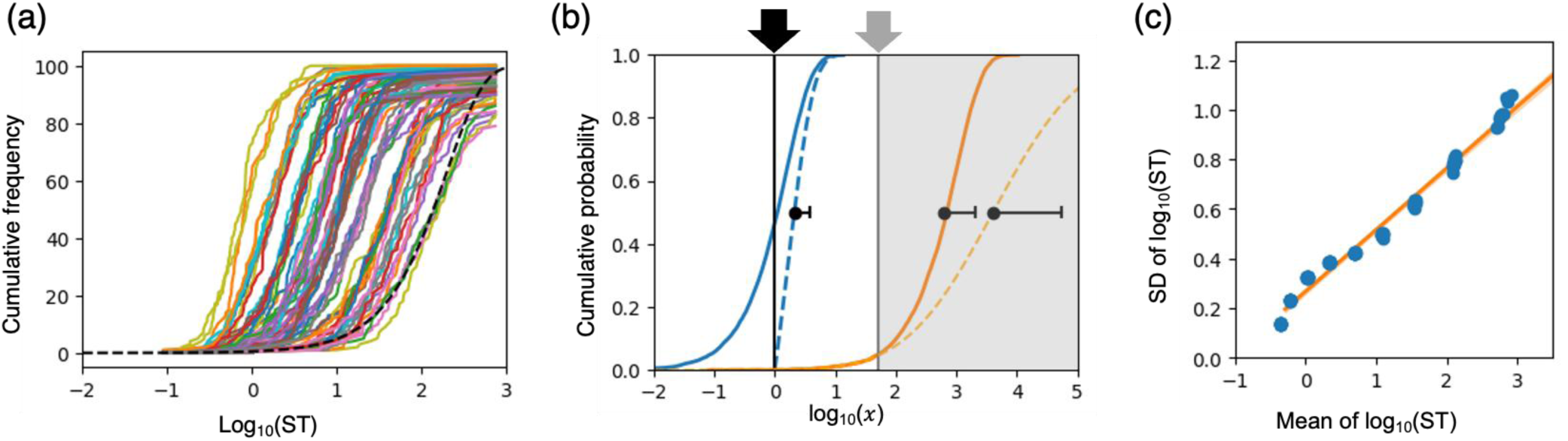
Mechanistic origin of the observed mean-SD correlation. **a** Simulation results across all images using the modified model in which the exponential term was replaced with a Boolean gating rule. The black dashed line indicates a pure exponential distribution. **b** Schematic illustration of two effects: a non-search time (black arrow) and a time-on-task limit (gray arrow). Cumulative distribution functions (CDFs) are shown for samples generated from pure exponential distributions with λ = 10^-0.2^ (blue) and 10^-3^ (orange) as representative examples; these two CDFs have identical shapes and differ only by a horizontal shift. The blue dashed line shows the result of adding a constant non-search time to ST, which narrows the fitted distribution (filled circles indicate the mean, and one-sided bars indicate the SD). The orange dashed line shows the result of fitting a lognormal distribution using only the data up to the time-on-task limit, illustrating that truncation can increase both the fitted mean and SD. **c** Relationship between the mean and SD obtained from lognormal fits after applying the two effects to pure exponential samples. The rate parameter λ was varied in equal logarithmic steps across nine values from 10^-0.8^ to 10^2.5^. For each λ, ten datasets were generated, each consisting of nine sets of 10000 exponentially distributed random samples. After applying the two effects, a lognormal distribution was fitted to each dataset. The resulting mean and standard deviation on the log scale are shown as a scatter plot. The slope, intercept, and R² were 0.25, 0.27, and 0.98, respectively.

For an exponential distribution, the SD in the log domain should be constant 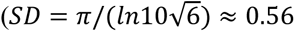 for base-10 logarithms). Nevertheless, when we fitted the resulting data with a lognormal distribution, we observed a proportional relationship between the fitted mean and SD. We identified two factors that can account for this proportionality in the lognormal fit: the presence of a non-search time and a time-on-task limit. Both of these factors can also arise in the previous human experiments (Murata et al., 2014).

First, adding a non-search time to ST compresses the left tail of the exponential distribution. As a consequence, lognormal fitting yields a smaller SD (Figure 4b). Converting RT to ST partially corrects for this delay, but does not eliminate it completely. This residual delay has a larger relative impact when RT is small, that is, for images with smaller *N_M_* values. As a result, SD is reduced more strongly for smaller-*N_M_* images, inducing an apparent proportionality between mean and SD across images in the low-*N_M_* range, where the fitted SD falls below 0.56. Consistent with this account, when we increased the non-search-time correction constant used in the RT-to-ST transformation, the correlation in the low-*N_M_* range disappeared (Online Resource 2).

Second, a time-on-task limit degrades the quality of lognormal fitting, particularly for images with large *N_M_* values. For such images, the cumulative frequency at the time-on-task limit remains below one half, meaning that the fitted median (the point at which the cumulative frequency reaches one half) must be obtained by extrapolation (Figure 4b). Therefore, when fitting variability yields a larger fitted mean, it tends to be accompanied by a larger fitted SD. This effect induces an apparent proportionality between mean and SD for images with larger *N_M_* values. Indeed, when we varied the mean parameter of a pure exponential distribution and applied these two effects, we reproduced an approximately proportional relationship between mean and SD similar to that observed in Figure 3a (Figure 4c).

Taken together, even though the distribution of pure search times follows an exponential distribution, applying the same processing steps used in the human experiments can yield a lognormal-like distribution with a proportional relationship between mean and SD, as observed in the human data. Nevertheless, because human experiments have reported stable individual differences across participants (Murata et al., 2014), we do not rule out participant-related sources of lognormality or other contributing factors. However, even in the absence of such factors, the lognormal-like distributions and the mean-SD proportionality reported in the human experiments could arise from the mechanisms described above.

## Conclusion

In this study, we proposed a model for generating STs in Mooney image recognition, a widely used paradigm for visual insight. The model takes the same input images used in human experiments and produces STs that exhibit the same statistical properties as those reported in the human data (Murata et al., 2014). It consists of two components: a DNN module that extracts perceptual visual features and an RNN-based chaotic search module that supplements missing features required for recognition. To the best of our knowledge, this is the first study to model human ST statistics in Mooney image recognition using ANNs for both the perceptual front end and the search process underlying recognition.

In particular, previous models based on random search (Hosoda et al., 2023) treat trial-to-trial variability as externally injected randomness and therefore provide limited leverage for mechanistically modeling stable individual differences or experience-dependent changes. In contrast, the model proposed here reproduces key statistical signatures of the human data while offering a neural-dynamical substrate in which such effects could, in principle, be modeled through synaptic plasticity. For example, human experiments have reported that participants who are fast at recognizing one image also tend to be fast at recognizing other images (Murata et al., 2014); such differences could potentially be represented as differences in neural network parameters. This mechanistic capability constitutes a key advantage of our approach and provides a concrete step toward a mechanistic understanding and neural implementations of insight-related recognition.

## Acknowledgements

We are grateful to the cluster computing team for their expert support. This study was supported in part by MIC under a grant entitled “R&D of ICT Priority Technology (JPMI00316)” and JSPS KAKENHI Grant Number JP24K15188 and JP25H01365.

## Author contributions

MK, KK, and KH designed the study and developed the concept. TM, MI, IO, TY, KK and KH provided supervision for the study. YM, SS, KK, and KH designed the model architecture. MK and KH carried out the computational experiments. MK and KH drafted the manuscript. All authors contributed critical revisions and approved the final manuscript for publication.

## Supplementary Information

**Online Resource 1.**
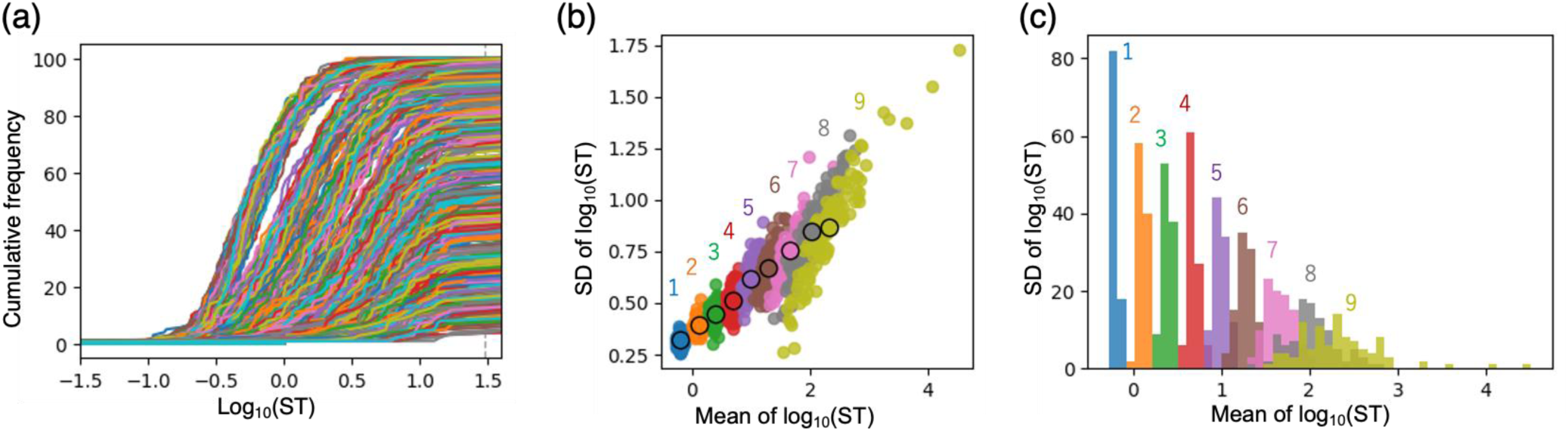
Statistical properties of search times (STs) across images for a large image set. The virtual images were generated by randomly sampling the visual feature vector ***h*** and the correct weight vector *w_c_*. For each value of *N_M_* from 1 to 9, 100 images were generated, resulting in a total of 900 images used in the simulation. **a**. Simulation results over the entire time course. Each line represents a single input image, and the vertical axis indicates the number of simulated participants who had completed the search by that time point. To approximate the time-on-task limit in human experiments, the cutoff time was set to equal to the value used for the main results. This cutoff is shown by the black dashed line. **b.** Relationship between the mean and standard deviation (SD) of log-transformed STs across images. Each point represents one of the 900 images. Colors indicate different values of *N_M_*, and the numbers represent the corresponding values of *N_M_*. Black-outlined circles indicate the mean values within each *N_M_* cluster. A proportional relationship between the mean and standard deviation is also observed among these cluster means. **c**. Histogram of the mean log-STs across images. Numbers indicate the corresponding values of *N_M_*. The spacing between *N_M_* clusters are approximately constant, indicating preserved discreteness. The apparent broadening of the cluster distributions with increasing *N_M_* is attributable to reduced estimation accuracy resulting from fitting the relationship between log_10_(ST) and cumulative frequency (a) for images with limited cumulative data. Therefore, this broadening does not reflect a substantive issue.

**Online Resource 2.**
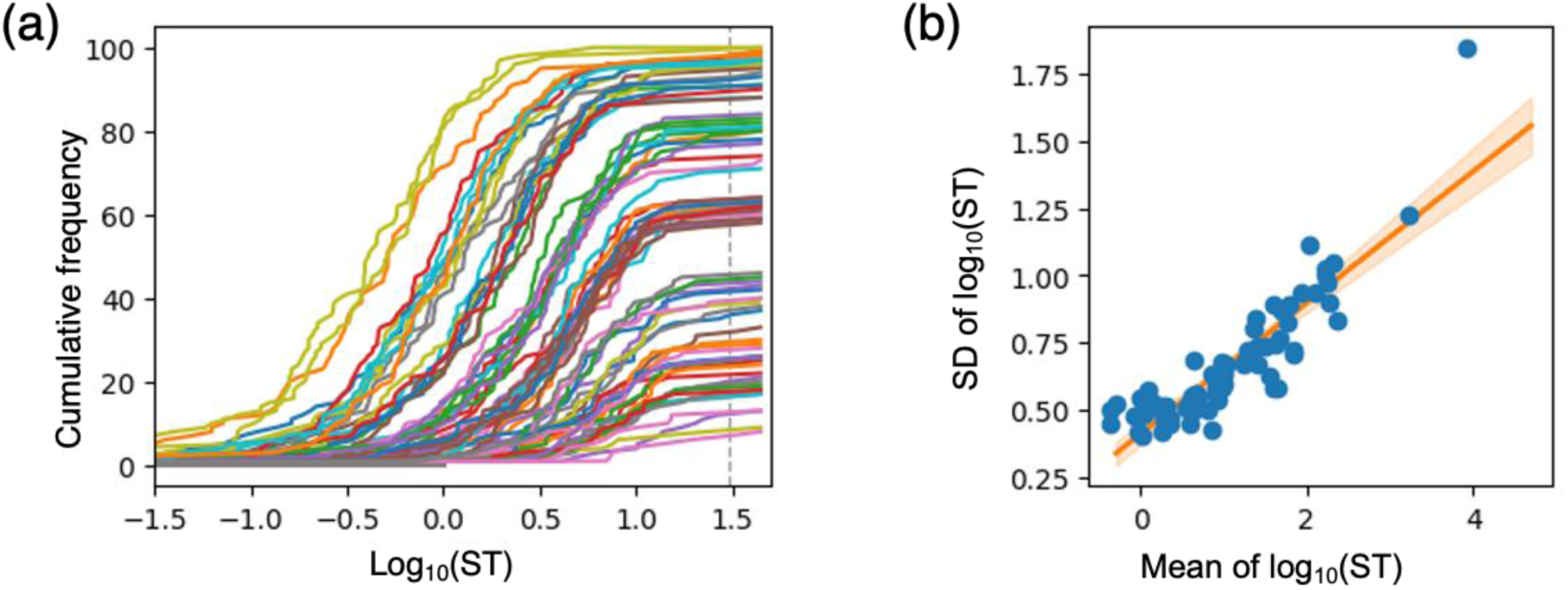
Statistical properties of fitted STs across images with an increased RT-to-ST conversion. In this condition, STs were computed by subtracting 2.5 from the RTs, instead of subtracting the minimum RT for *N_M_* = 0 (2.44) **a.** Simulation results over the entire time course. Each line represents a single input image, and the vertical axis indicates the number of simulated participants who had completed the search by that time point. To approximate the time-on-task limit used in human experiments, we defined the time at which the last participant finished searching the most difficult images as the 30-s cutoff. **b.** Relationship between the mean and standard deviation (SD) of log-transformed STs across images. Each point represents one of the 90 images. The line shows the result of linear regression, with the shaded area indicating the 95% confidence interval (CI). The slope and intercept of the regression line were 0.24 and 0.41, respectively, with a Pearson correlation coefficient of R = 0.89. Increasing the RT-to-ST conversion offset eliminated the proportional relationship observed at small *N_M_* values in the rising phase.

